# Binding site localization on non-homogeneous cell surfaces using topological image averaging

**DOI:** 10.1101/2020.11.03.366476

**Authors:** Vibha Kumra Ahnlide, Johannes Kumra Ahnlide, Jason P. Beech, Pontus Nordenfelt

## Abstract

Antibody binding to cell surface proteins plays a crucial role in immunity and the location of an epitope can altogether determine the immunological outcome of a host-target interaction. Techniques available today for epitope identification are costly, time-consuming, and unsuited for high-throughput analysis. Fast and efficient screening of epitope location can be useful for the development of therapeutic monoclonal antibodies and vaccines. In the present work, we have developed a method for imaging-based localization of binding sites on cellular surface proteins. The cellular morphology typically varies, and antibodies often bind in a non-homogenous manner, making traditional particle-averaging strategies challenging for accurate native antibody localization. Nanometer-scale resolution is achieved through localization in one dimension, namely the distance from a bound ligand to a reference surface, by using topological image averaging. Our results show that this method is well suited for antibody binding site measurements on native cell surface morphology.

## Introduction

The location of protein binding sites on cellular surfaces can have wide-ranging implications for various cellular processes, such as immune signaling, cell adhesion, cell migration, and phagocytosis. Antibody binding to pathogen surface proteins plays a crucial role in immunity (1), and epitope localization affects diverse immunological outcomes, such as antibody neutralizing ability (2) (3) (4) (5) (6) or autoreactivity (5) (7). Even though monoclonal antibody (mAb) treatments have proven to be successful for a wide spectrum of disease, there are very few monoclonal antibodies clinically available for the treatment of bacterial infections (8). Developing therapeutic mAbs against infections may require large scale screening of highly conserved and functional epitopes (9). Antibody epitopes are typically identified using crystallography, mutagenesis, or crosslinking coupled mass spectrometry (10). As of today, these methods are costly, time-consuming, and unsuited for high-throughput analysis. Fast and efficient screening of epitope locations can be useful for the development of therapeutic mAbs and vaccines.

While there exist averaging methods for nanometer-scale localization on microscopy data, these do not work for binding sites on proteins with a non-homogenous expression on the cellular surface. Moreover, they are designed for identical (11) (12) or spherical (13) particles and are thereby not fitted for unaltered cells due to the existing biological variation of surface morphology.

In the present work, we have developed an imaging-based method for localization of binding sites with nanometer-scale precision. The location of a binding site is determined by calculating the mean distance between the ligand channel and a reference signal channel, using the averaged fluorescent signal along a cell contour. We apply this method to determine known binding site locales on M protein that is unevenly expressed on the bacterial surface of *S. pyogenes.* Our presented results with structured illumination microscopy (SIM) data achieve a ~5 nm precision. Additionally, our method yields viable results also when we perform site localization on wide-field images with and without deconvolution. To sum up, our site localization method enables rapid determination of binding sites, without the need for synthetic modification of cell surface morphology.

## Results

### Principles of site localization method

Binding site localization is based on resolving the axial distance between bound ligands and a reference surface. This is done by implementing an averaging method on high-resolution images. With this method, nanometer-scale precision is achieved by dimension reduction of surface data within an image plane.

A site localization measurement on antibody-coated bacteria is exemplified in Fig. 1. Fluorescently conjugated wheat germ agglutinin (WGA) that binds bacterial peptidoglycan is used here for the reference region and antibodies that bind to bacterial M protein are labeled with another fluorescent dye for imaging in a separate channel (Fig. 1a). The example shows how our method handles the uneven expression of M protein on bacterial surfaces (Fig. 1b). The pipeline for the site localization method is shown in Fig. 1b. For each image, bacteria are identified in the reference channel by fitting circles to an edge-detection-processed image. Each circle is isolated as a slightly larger mask and the data in the circle is polar transformed. An alignment of the reference position is performed by identifying peaks to each intensity profile. This way, the axial positions are normalized to the actual surface signal. The spatially corresponding data in the antibody channel is processed in the same manner and the distance between the antibody and WGA signal is resolved through the averaging of multiple acquisitions of the same field of view. Altogether a relative binding site is determined by resolving the distance between the reference and antibody channel using the cumulative measurement from multiple repeated images.

**Fig. 1.**
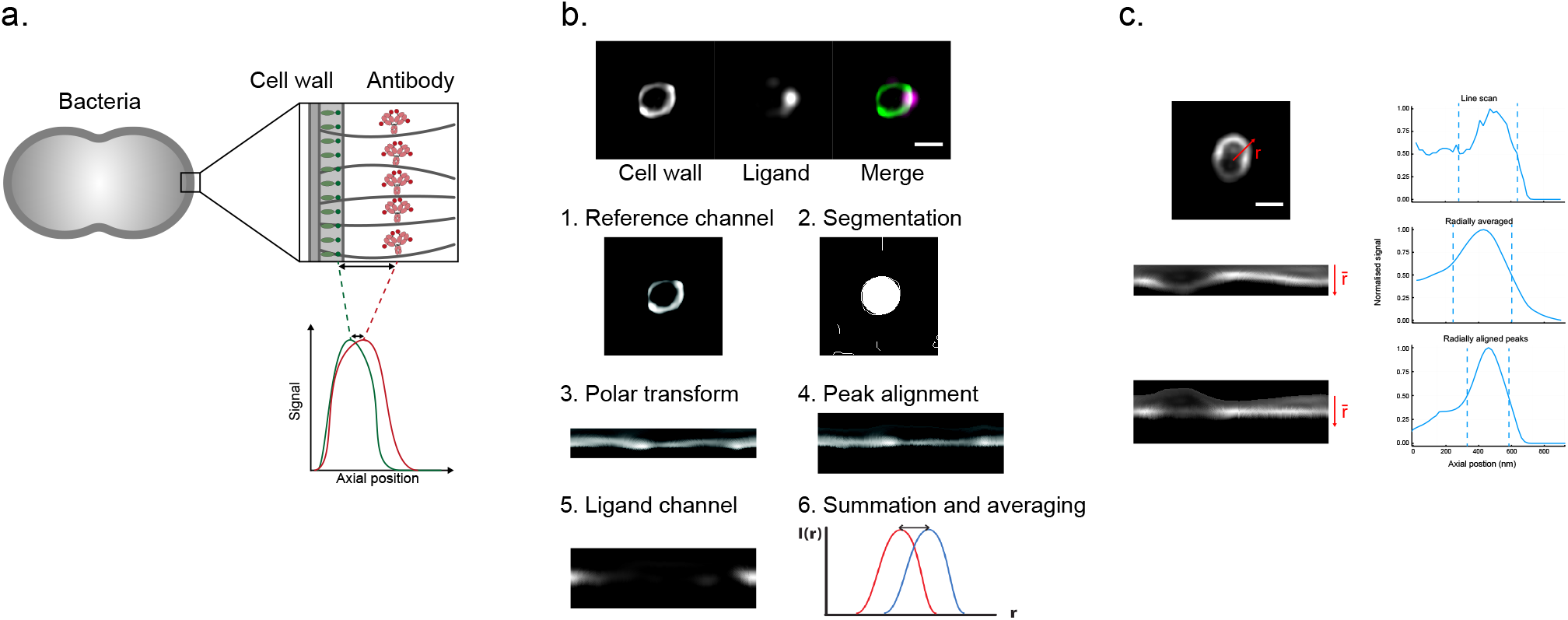
Binding site localization is based on resolving the distance between bound ligands and a reference surface. **a. Site localization on bacterial surfaces** The illustration exemplifies a measurement with antibody-coated bacteria. Antibodies with red fluorescent dyes are bound to bacterial surface proteins. A bacterial cell wall binding protein is labeled with a green fluorescent dye. High-resolution images of single bacteria are acquired in the focal plane and the fluorescent peak signal is averaged along the bacterial contour. The distance *d* is determined by resolving the difference between the fluorescent signal peaks. **b. Pipeline for the site localization method** Structured Illumination Microscopy (SIM) images are shown at the top. *S. pyogenes* strain SF370 has been fixed, stained with AlexaFluor488-conjugated wheat germ agglutinin (WGA), and coated with antibody Fc fragments. The antibody fragments were stained with Fc specific AlexaFluor647-conjugated F(ab’)_2_ fragments. The scale bar is 500 nm. The aim is to locate the antibody binding site by calculating the mean distance between the antibody channel and a reference signal channel. 1. A raw image in the reference channel is shown. 2. Bacteria are identified by fitting circles to an edge-detection-processed image. 3. The data in each circle mask is isolated and transformed to polar coordinates. 4. An alignment of the reference position is performed by identifying peaks at each radial position. 5. The spatially corresponding data in the antibody channel is extracted. 6. The radially aligned peaks are then averaged. The peak distance between these intensity profiles should then correspond to the distance between the binding site and the bacterial peptidoglycan layer. **c. Representation of improved SNR by peak alignment for non-spherical particles** Intensity profiles (right) for the reference channel are shown together with their respective image data (left). The signal along a single line for an oval-shaped bacteria is shown at the top. The intensity profile of a radial average is shown in the middle. An improved SNR is seen as a peak alignment is performed (bottom).

A strength of the method is that it can be used for particles with a non-spherical topology. It is evident that the signal precision is improved as the distortion of an ovoid bacteria is accounted for through axial normalization in polar coordinates (Fig. 1c).

We tested and compared our method to an existing method for height measurement on cellular surfaces, CSOP (13) (see Fig. S2). CSOP failed to detect accurate circles in the ligand channel, owing to the non-homogenous expression of M protein on the cellular surface. Moreover, as CSOP does not account for non-spherical topology this method is not as suited for site localization on unaltered bacterial surfaces. This is likely the reason results with CSOP give lower precision than our site localization method for bacteria with uniform labeling.

### Validation of site localization method by measurement of binding sites on bacterial surface protein

Site localization measurements that were performed for ligands on bacterial M protein agree well with previously reported data. The antibody Fc binding site on M protein is located at the S region (14) and a non-specific monoclonal IgG antibody (Xolair) is used for site localization of this binding. Additionally, a measurement is carried out for an M protein specific monoclonal antibody, Ab49 (Bahnan et al 2020, manuscript), with an epitope located in the B3-S region, i.e. slightly further along the IgGFc binding region. For determining binding to the far end of M protein, we used fibrinogen that has two binding sites at the B1 and B2 region (15). We thus expect the binding sites to be arranged in accordance with the schematic shown in Fig. 2c. *S. pyogenes* strain SF370 was stained with AlexaFluor488-conjugated WGA and coated with the Fc fragment of Xolair, F(ab’)2 fragment of Ab49 or AlexaFluor647-conjugated fibrinogen. The antibody fragments were stained with either Fc or Fab specific AlexaFluor647-conjugated F(ab’)_2_ fragments. Images of single bacteria were acquired using an N-SIM microscope. The mean resolved distance is 36.7±4.7nm for the Xolair Fc binding site, 49.0 ± 5.9nm for Ab49 Fab binding site and 83.0 ± 13.3 nm for the two fibrinogen binding sites. These results are consistent with the arrangement of the binding sites as reported in the literature (14) (15) (Bahnan et al 2020, manuscript).

**Fig. 2.**
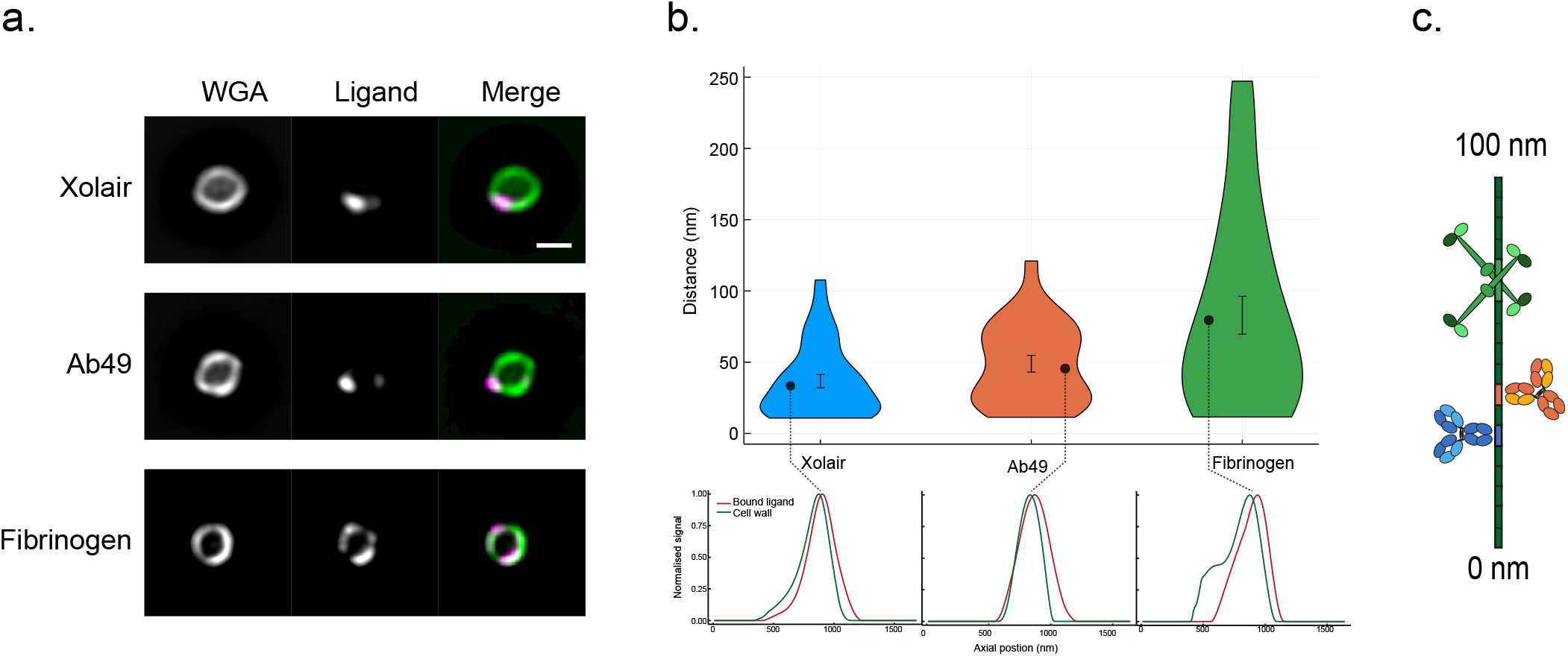
Site localization measurements of bound ligands on bacterial surface protein. a. Representative SIM images of ligand coated bacteria. *S. pyogenes* strain SF370 has been fixed, stained with AlexaFluor488-conjugated wheat germ agglutinin (WGA) and coated with the Fc fragment of Xolair (top), F(ab’)_2_ of Ab49 (middle) or AlexaFluor647-conjugated fibrinogen. The antibody fragments were stained with either Fc or Fab specific AlexaFluor647-conjugated F(ab’)_2_ fragments. Scale bar is 500 nm. **b. Binding site measurements of antibodies and fibrinogen on bacterial M protein** The measured distances are shown in a violinplot. The mean resolved distance is 36.7 ± 4.7nm (n = 55) for the Xolair Fc binding site, 49.0 ± 5.9 nm *(n* = 37) for Ab49 Fab binding site and 83.0 ± 13.3 nm *(n* = 42) for two fibrinogen binding sites. Error bars indicate standard error of mean. An intensity profile from a single bacteria is shown for each of the ligands below the violinplot. **c. A schematic illustrating the determined binding sites on M protein.** The relative binding sites agree well with reported data in the literature.

**Fig. 3.**
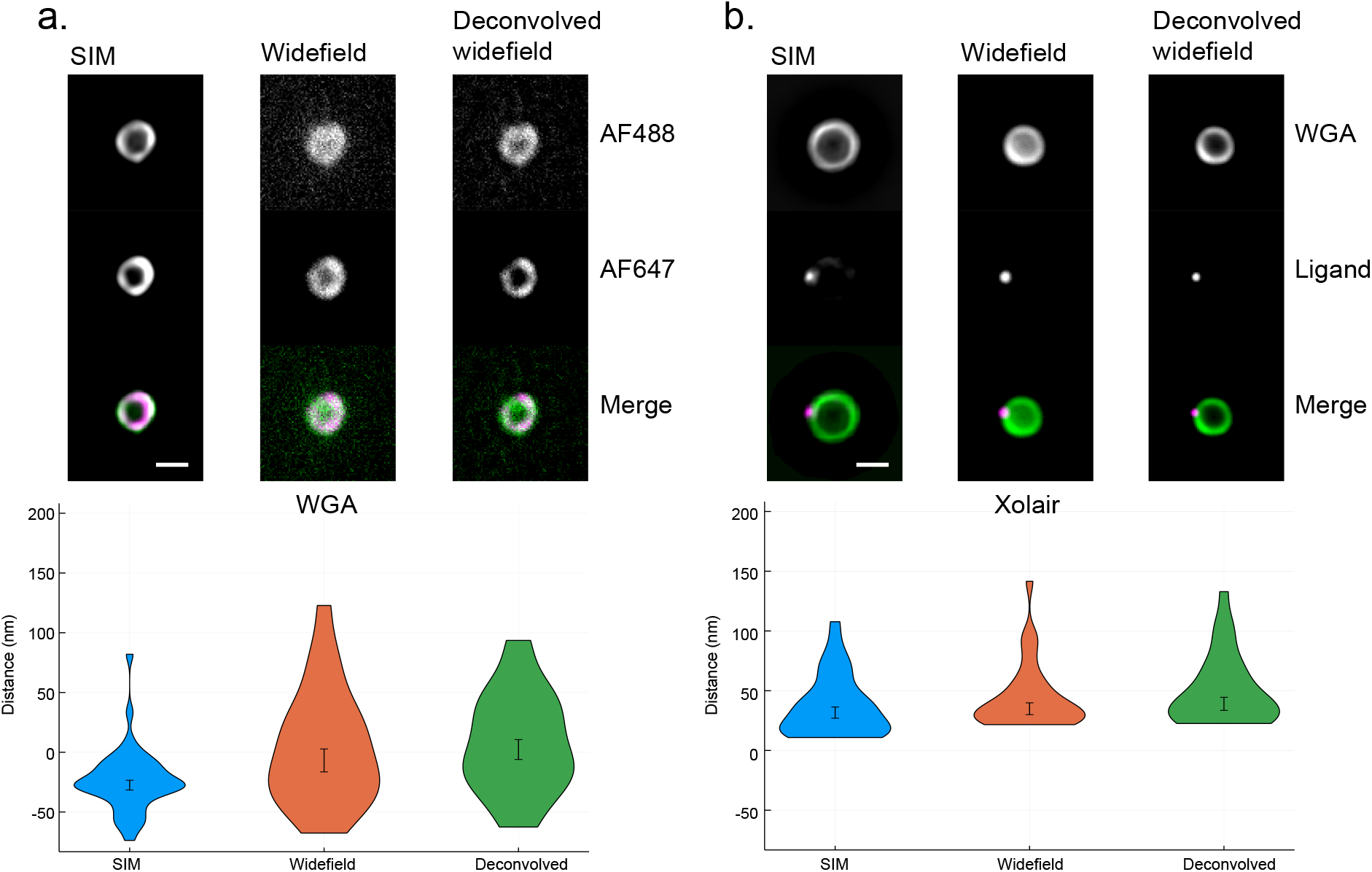
A comparison of site localization measurements using SIM, widefield, and deconvolved widefield images. **a. Site localization performed for two-channel cell wall signal** Representative images are shown at the top. *S. pyogenes* strain SF370 has been fixed and stained with AlexaFluor488- and AlexaFluor647-conjugated WGA. Scale bar is 500 nm. SIM and widefield images are acquired of the same set of bacteria (n =25). The mean resolved distance between the two channels is −27.0 ± 4.1 nm with SIM data, −6.8 ± 9.6 nm with widefield data and 2.3 ± 8.3 nm with deconvolved widefield data. Error bars indicate standard error of mean. **b. Site localization measurement of Fc binding site yields similar results with widefield dataset** Representative images are shown at the top. *S. pyogenes* strain SF370 has been fixed, stained with AlexaFluor488-conjugated WGA, and coated with antibody Fc fragments. The antibody fragments were stained with Fc specific AlexaFluor647-conjugated F(ab’)_2_ fragments. Scale bar is 500 nm. SIM and widefield images are acquired of two distinct sets of bacteria. Measured distances are shown in the violin plot. The mean resolved distance for the Xolair Fc binding site is 30.7± 4.7nm (n =30) with SIM data, 35.0 ± 5.0 nm (n =31) with widefield data and 39.0 ± 5.5 nm (n =29) with deconvolved widefield data. Error bars indicate standard error of mean.

### Site localization measurements using widefield and deconvolved images

By performing site localization measurements using widefield images, we show that our method yields high precision even with conventional microscopy data. Moreover, deconvolution of the widefield images may give a slight increase in precision. Widefield images were acquired with the same configurations and optical system as the SIM images (see previous section), with the exception of the light source used being a LED-based Lumencor Spectra X. Comparison of site localization measurements using SIM and widefield were performed on two sets of samples; *S. pyogenes* with two channel cell wall signal and Xolair-coated *S. pyogenes*. For the first set, bacteria were stained with both AlexaFluor488- and AlexaFluor647-conjugated WGA Fig. 2a. In this case, SIM and widefield images were acquired of the same set of bacteria. The mean resolved distance between the two channels is −27.0 ± 4.1 nm with SIM data, −6.8 ± 9.6nm with widefield data and 2.3 ± 8.3 nm with deconvolved widefield data. The difference in precision for double-stained WGA data is likely because of decreased SNR in widefield images due to photobleaching by prior SIM imaging. For the Xolair samples, SIM and wide-field images were acquired of two distinct sets of bacteria. The mean resolved distance for the Xolair Fc binding site is 31.7 ± 4.7 nm with SIM data, 35.0 ± 5.0 nm with widefield data and 39.0 ± 5.5 nm with deconvolved widefield data. This indicates that the site localization method performs well with widefield images as long as there is sufficient signal.

## Discussion

The locations of protein binding sites on cellular surfaces are important for a wide range of cellular processes. In particular, how and where an antibody binds to pathogen surface structures is critical for the outcome of the hostpathogen interaction. This is especially relevant for bacteria as they are known to target antibodies in many different ways (16) (17). Knowing the location of the epitope greatly aids both mechanistic understanding and facilitates potential therapeutic development.

We have developed a method for imaging-based localization of binding sites on cellular surface proteins with nanometer-scale precision. The practical resolution of super-resolution techniques such as iPALM, STORM and STED is ~10nm. Here, a ~5 nm precision is achieved with diffraction-limited microscopy images by dimension reduction of surface data within an image plane. Existing imaging based averaging methods are designed for identical or spherical particles and therefore not suited for site localization on native cell surface morphology. This is made possible with our method through normalization of the axial positions of a reference surface.

We have applied our site localization method to determine known binding site locales on M protein that is unevenly expressed on the bacterial surface of *S. pyogenes.* Our presented results show that this method is suited even for cells with non-spherical topology. Finally, we show that our method yields viable results with conventional microscopy and we demonstrate that the precision may be increased by deconvolution of widefield images. Improvements, such as the use of site-specific fluorescent conjugation may increase the measurement precision. We believe this method may be useful for rapid screening of epitope locales and the implementation, written in Julia, is provided on GitHub (VibhaKumra/SiteLocalization/tree/v0.1.0).

## Materials and Methods

### Bacterial culturing conditions

*S. pyogenes* strain SF370 wildtype and ΔM mutant (18) were cultured overnight in THY medium (Todd Hewitt Broth; Bacto; BD, complemented with 0.2% (w/v) yeast) at 37 °C in an atmosphere supplemented with 5% CO_2_. Strain SF370 expresses M1 protein on its surface and is available through the American Type Culture Collection (ATCC 700294) (19). The bacteria were harvested at early log phase and washed twice with PBS.

### Opsonisation of bacteria

Xolair (Omalizumab, Novartis) is a humanised monoclonal IgG that is IgE-specific, and thus only binds to M protein via Fc. Ab49 is an M protein specific antibody (Bahnan et al, manuscript). Both antibodies were treated with IdeS (Hansa BioPharma) (20), an enzyme that cleaves IgG at the hinge region, separating the F(ab’)2 from the Fc. The fibrinogen used here was isolated from human plasma and conjugated with AlexaFluor-647 (Invitrogen™).

### Fixation and staining

Bacteria were sonicated (VialTweeter; Hielscher) for 0.5 minutes to separate any aggregates and incubated fixed in 4% paraformaldehyde for 5 minutes on ice. The bacteria were thereafter washed with PBS twice (10000g, 2min). SF370 wild type was stained with Alexa Fluor 488-conjugated WGA. Bacteria were incubated with IdeS-cleaved Xolair, Ab49 or AlexaFluor647-conjugated Fibrinogen (Invitrogen™). The antibody samples were stained with fluorescently labeled IgGFab or IgGFc specific F(ab’)2 fragments (AlexaFluor647-conjugated anti-human IgGFc or IgGFab; Jackson ImmunoResearch Laboratory). Samples were set on glass slides using Prolong™ Gold Antifade Mountant with No. 1.5 coverslips.

### SIM image acquisition

Images of single bacteria were acquired using an N-SIM microscope with LU-NV laser unit, CFI SR HP Apochromat TIRF 100X Oil objective (N.A. 1.49) and an additional 1.5x magnification. The camera used was ORCA-Flash 4.0 sCMOS camera (Hamamatsu Photonics K.K.) and the images were reconstructed with Nikon’s SIM software on NIS-Elements Ar (NIS-A 6D and N-SIM Analysis). Fluorescent beads (100 nm) were imaged to measure and correct for chromatic aberration, as well as for the N-SIM grating alignment. Single bacteria were manually identified and imaged with 488 and 640 nm lasers in time series with 20 frames.

### Microscope calibrations

TetraSpeck™ 0.1 μm fluorescent microspheres are mounted on No. 1.5 coverslips using Prolong™ Gold Antifade Mountant, in the same manner as the bacterial samples. These beads are used for the objective collar correction, SIM grating alignment and measurement of SIM and widefield PSF. Images of the beads were acquired and chromatic aberration was corrected for by performing image registration (Holylab/RegisterQD.jl) and applying the found transform to all images.

### Site localization analysis

The analysis pipeline was implemented in Julia and is available on GitHub. A Circle Hough Transform (21) and Canny Edge Detection (22) were used for circle fitting and edge detection, respectively. A polar transform of the found circle was performed on a bicubic interpolation of the image. The alignment of the peak intensity was performed by identifying a peak maximum using a sliding average. A low pass filtration was performed on the alignment to smoothen out potential high-frequency noise. To avoid attaining out-of-focus peaks in the ligand channel, peak identification was performed beyond the reference region. Number of bacteria together with mean and SEM are given in the figure captions above. The widefield images were deconvolved using the Richardsson-Lucy algorithm (23) (24) in 10 iterations.

## ACKNOWLEDGEMENTS

Jonas Tegenfeldt and Elke Hebisch for support during early phase of the method development.

## Supplementary Note 1: Double cell wall stain control

**Fig. S1.**
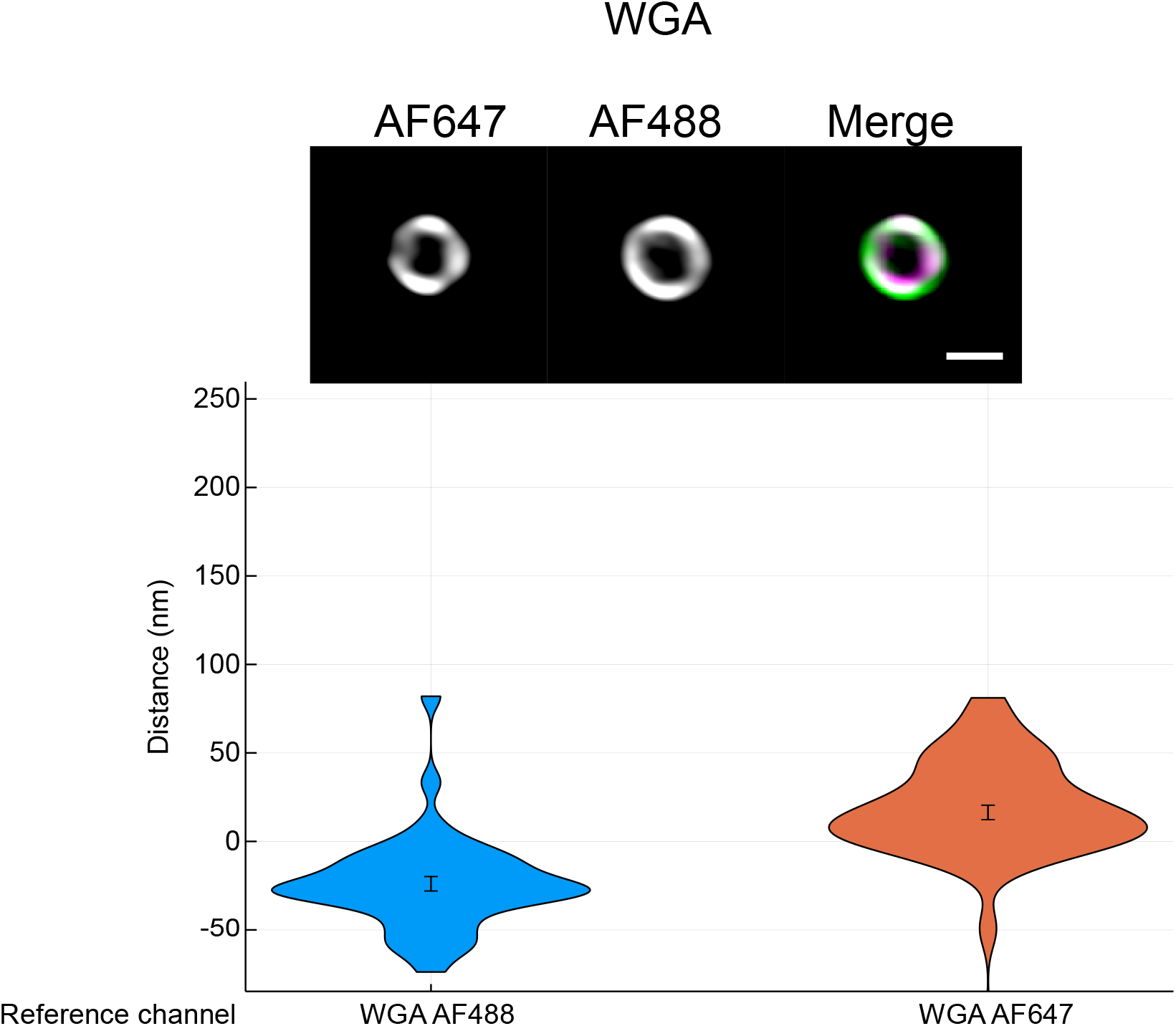
Control of site localization method using double stain wall control. Representative images are shown at the top. *S. pyogenes* strain SF370 has been fixed and stained with AlexaFluor488- and AlexaFluor647-conjugated WGA. Scale bar is 500 nm. The violin plots show the resolved distance between the two cell wall signals. The mean resolved distance between the two channels is −23.9 ± 4.1 nm with AlexaFluor647 as reference and 16.4 ± 5.0 nm with AlexaFluor488 as reference. A mirroring of the results is expected as the reference and target channels are flipped and this is evident from the shape of the violin plots.

## Supplementary Note 2: Comparison of Site localization and CSOP

**Fig. S2.**
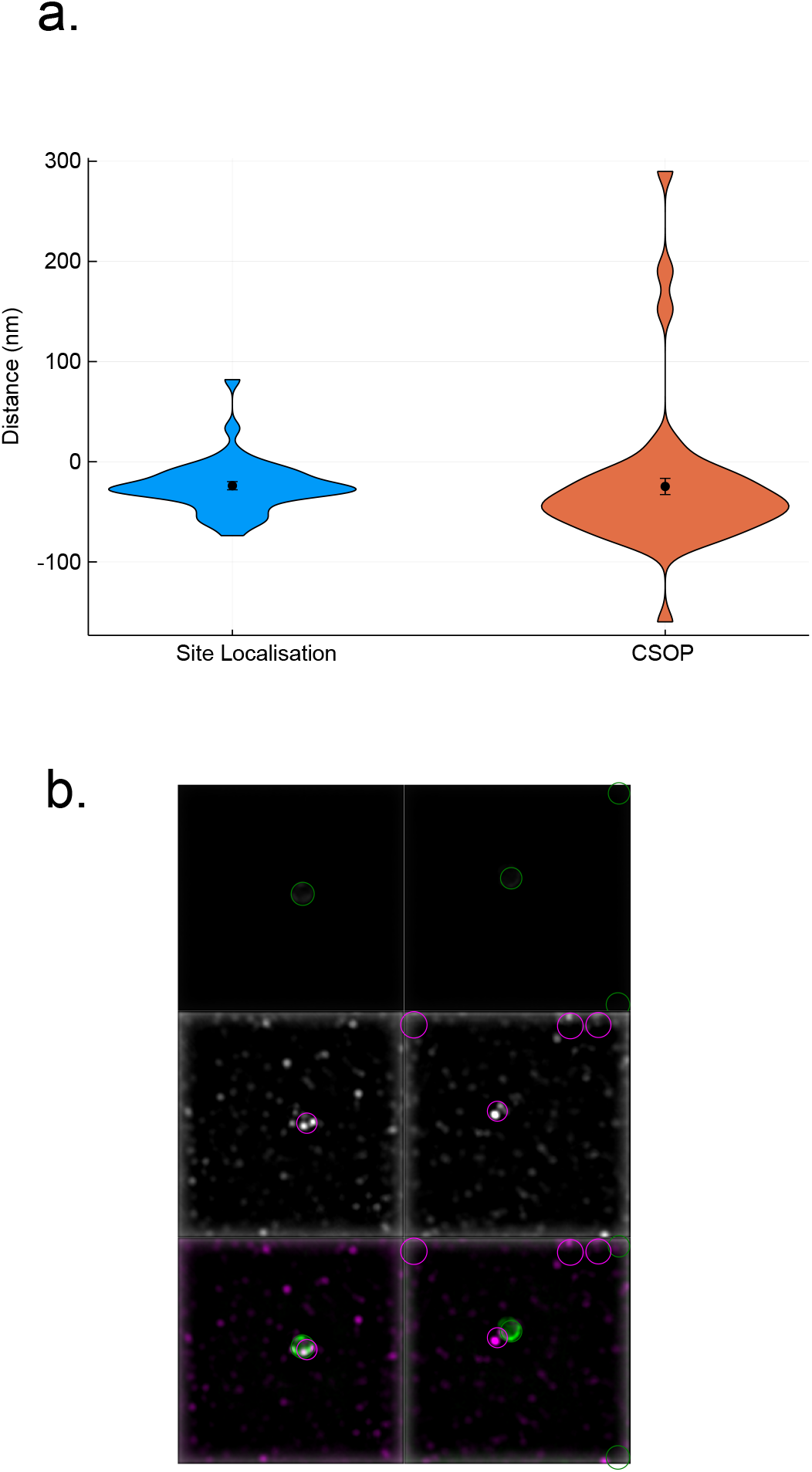
Distance measurements with site localization and CSOP. **a. Distance determination using CSOP and site localization** The measurements are performed for the same set of data. *S. pyogenes* strain SF370 has been fixed and stained with AlexaFluor488- and AlexaFluor647-conjugated WGA. The mean resolved distance between the two channels is −23.9 ± 4.lnm with site localization and −24.7 ± 8.lnm with CSOP. The larger variation in the result with CSOP is likely due to the fact that the non-spherical topology of *S. pyogenes* is not accounted for with this method. **b. Examples illustrating difficulty using CSOP for localization of binding on proteins with patchy expression on the cellular surface** The top images are the cell wall channel, the middle images the ligand channel and the bottom ones are a merge of the two. Images analysed using smson-ucb/CSOP. CSOP performs circle segmentation in both the reference and target channel. The patchy protein expression has given rise to faulty circle detection.

